# Differences in energy storage in sympatric salmonid morphs with contrasting lifestyles

**DOI:** 10.1101/2024.08.14.607877

**Authors:** Evgeny V. Esin, Grigorii N. Markevich, Elena V. Shulgina, Yulia A. Baskakova, Roman V. Artemov, Fedor N. Shkil

**Affiliations:** A.N. Severtsov Institute of Ecology and Evolution RAS; Moscow, Russian Federation; Vitus Bering Kamchatka State University, Petropavlovsk-Kamchatsky, Russian Federation; Russian Federal Research Institute of Fisheries and Oceanography; Moscow, Russian Federation

**Keywords:** Biochemical parameters, Lipids, Glycogen, Proteins, Metabolism, Charr, Kronotskoe

## Abstract

Although physiology ensures homeostasis and fitness in a particular environment, and ecological shifts cannot be realized without physiological changes, metabolic transformations during animal adaptive radiations still remain unexplored. We present a study of energy reserve storage in the salmonid assemblage inhabiting a cold-water Lake Kronotskoe. This assemblage diversified from *Salvelinus malma* and includes eight distinct ecomorphs with contrasting lifestyles and trophic specializations. We hypothesized that ecomorphs differ in energy storage and expenditure, and that their metabolic phenotypes should be among the primary targets of natural selection. To test this hypothesis, we compared the stored amount and ratio of carbohydrates, lipids, circulating peptides that supply the citric acid cycle, as well as proxy indicators of metabolic rate, the blood levels of plasma proteins (including albumin) and hemoglobin. Among ecomorphs, numerous significant differences in physiological parameters were found, closely related to the composition of food, the depth of habitat, and determined by internal factors, probably genetics. Each ecomorph has a specific metabolic phenotype corresponding to its tropho-ecological specialization and lifestyle. Metabolically advanced predators accumulate lipids; littoral insectivorous morphs grow slower and accumulate glycogen; amphipod feeders do not accumulate spare substances; the deepwater consumer of silt benthos differs in the most divergent physiological characteristics. We assume a specific selection on endocrine regulators of energy metabolism during the adaptive radiation of the assemblage, among which the most plausible candidates are thyroid hormones and leptin.

## INTRODUCTION

Adaptive radiation in fishes has been actively studied over the past decades, yielding a wealth of data revealing various factors and mechanisms underpinning morphological and ecological diversification. Since physiology ensures homeostasis and fitness under current external conditions (Jobling, 1994; Koedijk et al., 2010; Seebacher et al., 2015; Cloyed et al., 2019), ecological shifts and adaptations cannot be realized without physiological changes. Several studies have considered physiological adaptations in the intraspecific divergence of freshwater fishes (Proulx and Magnan, 2002; Goetz et al., 2013; Chavarie et al., 2016; Busarova 2017a; Esin et al., 2021), but their energetic basis remains largely unexplored. Studying the diversity of physiological phenotypes and their evolutionary dynamics should undoubtedly enrich our understanding of how ecological and morphological diversity in radiating fishes arises.

Here we present a study of the basic physiological characteristic – energy storage – in the ecomorph assemblage of salmonids inhabiting a cold-water dimictic lake (Lake Kronotskoe, North Asia). Energy metabolism is an inherent property of all life processes and a key factor in ecological adaptations (Brett and Groves, 1979; Hochachka and Somero, 2002; Van de Pol et al., 2017; Zwahlen et al., 2024). The composition and ratio of biopolymers involved in energy storage (carbohydrates, lipids, circulating peptides) underlie the survival and fitness in various environments (Goetz et al., 2013; Chavarie et al., 2016; Esin et al., 2021). Therefore, the study of metabolic energy processing in ecologically distinct morphs recently derived from the same ancestor is of great interest.

The Lake Kronotskoe ecomorph assemblage provides an excellent opportunity to study the transformation of energy metabolism during adaptive radiation. This assemblage diversified during the Holocene from the Dolly Varden charr, *Salvelinus malma* (Walbaum 1792), and consists of several reproductively isolated, morphologically distinct groups with contrasting lifestyles and trophic specializations (Markevich et al., 2018; Esin et al., 2020). Given that different diet and lifestyle require different physiology, we suggested that ecomorphs differ in energy storage and expenditure, and their metabolic phenotypes should be among the primary targets of natural selection.

To test this hypothesis, we compared the stored amount and ratio of biopolymers that supply the citric acid cycle in eight ecomorphs. We assessed the liver glycogen content as a stored source of glucose (Polakof et al., 2012) and the blood glucose level as an immediate energy reserve (Masanori and Shizunori, 1971). Muscle triacylglyceride content was analyzed as a stored source of fatty acids, and the ratio of different unbound (free) fatty acids was measured to evaluate sources of metabolic energy through β oxidation (Sargent et al., 2002; Tocher et al., 2003). We assessed the level of plasma proteins (including albumin), as a secondary source of oxaloacetate and related substances of the citric acid cycle (Andoh et al., 2007; Sawers, 2015; Falco et al., 2020), and since they are involved in the transport of various lipid, hormone and low-molecular compounds (Ronald and Taylor, 1977; Brett and Groves, 1979), as a proxy indicator of metabolic rate. We also measured the hemoglobin levels in the blood as an indicator of the intensity of oxygen transport and metabolic activity.

We assumed that the amounts and ratios of storage compounds would be related to the ecomorph’s lifestyle, type of feeding (diet) and depth of habitation. Ecomorphs that feed similarly should have a similar pattern of energy storage. The overall metabolic rate should be correlated with the hemoglobin level, and metabolically advanced ecomorphs may exhibit less correlation between tissue composition and ecological features due to their ability to process stored substances.

## MATERIAL AND METHODS

### Ecosystem description

Kronotskoe is a large (246 km^2^) and deep (up to 136 m) mountain lake situated on the west coast of the Kamchatka Peninsula (Fig. 1, incut). The ecosystem is cold-dimictic and oligotrophic; the lake depression consists of a littoral (≤ 12 m depth) that warms up to 15⁰С in summer, a slope and an extensive afotic profundal (> 30 m) with a constant temperature close to 4⁰С (own data). The lake is isolated from the downstream fish fauna, as there are unpassable rapids along the outflow river (Braitseva et al., 1995).

**Fig. 1.**
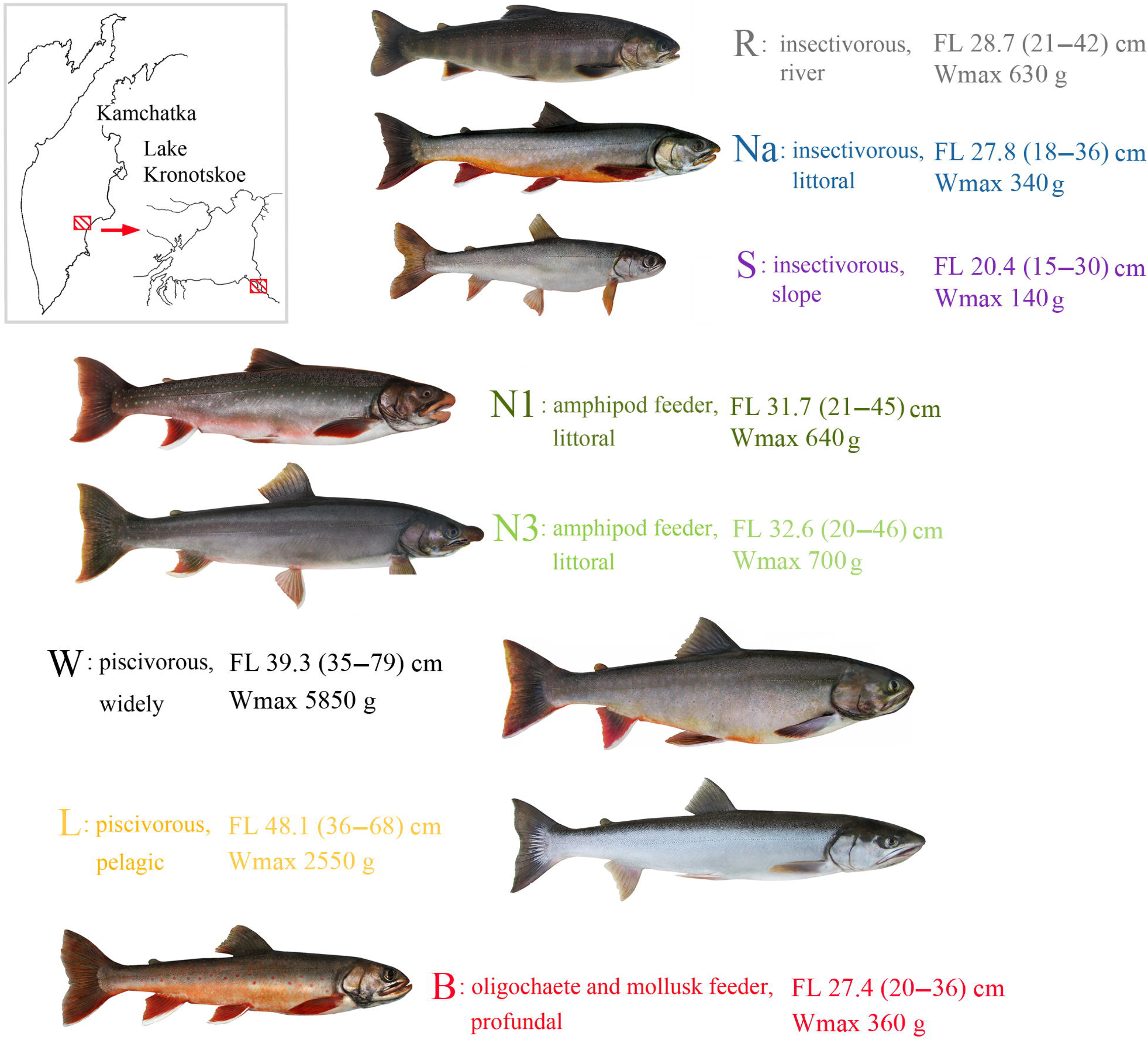
External appearance and sampling location of eight morphs of Lake Kronotskoe charr, key aspects of their diet and habitat partitioning, and parameters of body size in mature fish under analysis

Endemic charrs forage in the lake from June to early August and migrate to spawn at the specific remote sites in August – September (Markevich et al., 2021). Several morphologically specific groups with different feeding and habitat preferences could be differentiated in the ecosystem (Markevich et al., 2018; Esin et al., 2020). Littoral nosed charr (Na) and smallmouth charr (S) that inhabit the lake slope feed mainly on insect larvae and pupae (predominantly chironomids). Littoral charrs with lipped (N1, = N1g in our previous publications (see Busarova et al., 2017b)) and shovel-nosed (N3) morphotypes feed exclusively on amphipods (*Gammarus* spp.). Two large-sized morphs are mainly piscivorous: the most abundant widehead charr (W) and the pelagic longhead charr (L). The slow-growing profundal bigmouth charr (B) feeds on oligochaetes (Naididae) and bivalves (*Pisidium s*pp.) from the silt. The eighth morph (roundhead, R) inhabits the source of the outflowing river and feeds mainly on insect larvae, but also swallows juvenile fish running down from the lake (Fig. 1). The stability of the morphs’ dietary and habitat preferences was previously confirmed by their discrete isotopic niches (Esin et al., 2020). Analysis of the stomach contents and habitat distribution of fish sampled in this study is fully consistent with previous findings. Up to half of the mature fish skip the reproduction of the appropriate year and do not migrate to spawn, continuing to actively forage in the lake in August. These individuals do not develop sexual products and breeding dress.

### Material collection

We sampled charrs in littoral, slope, profundal habitats and in the water column, as well as in the source of the outflowing river in August 2022 using multi-panel gillnets (18-40 mm mesh size). To minimize injury and stress, gillnets were checked hourly, sampled fish were placed in the aerated container and immediately transferred to mesh cages (1.5 m^2^) installed in the water flow zone near the shore. Fish showing signs of reduced vital activity were rare and were removed from the cages. In order to standardize the data on the physiological condition of the charrs, only adults with no signs of breeding dress that had skipped the autumn spawning were sorted for further analysis. Later at autopsy, the fish were sexed and their status was verified on the basis of gonad weight ≤ 6% of body weight (additional criteria: a small proportion of oocytes in the stage of vitellogenesis in the ovaries, spermatocytes in the early stages of differentiation in the testes). From 16 to 20 such individuals of each morph were measured for body size (Fig. 1), kept in cages for 24 h and then processed following the standard protocol.

To ensure the charrs were in the same physiological condition, we measured blood glucose concentration using an automatic glucose-meter (Contour TS) prior to sampling. Blood glucose, which serves as an immediate energy reserve (Masanori and Shizunori, 1971), tends to fluctuate in response to feeding state. We observed similar glucose levels in all comparison groups (ranging from 5 to 15 nMol ml^-1^, with a mean of 8.98 nMol ml^-1^). There were no significant differences in this parameter (ANOVA F_7;122_ = 2.5 *p*=0.053), which can be attributed to the starvation state of all fish.

We did not consider variation in parasitic infestation due to the inaccessibility of rapid and effective treatment technology. Furthermore, specific parasite communities have been identified for the Lake Kronotskoe ecomorphs (Esin et al., 2020), which may force divergent patterns of physiological adaptation. Thus, disruption of the natural parasite load can have uncontrolled effects on the physiological state of the fish.

### Study protocol

Blood (2–4 ml) was collected from the caudal vessel using Vacuette serum tubes. Fish were not anaesthetized prior to blood sampling. The hemoglobin concentration was measured in 20 µl samples by the cyanmethemoglobin method (Gammon and Baker, 1977) with 5 ml of transforming reagent (Renam kit), and the remaining blood was centrifuged at 2900 g (Velocity 6u, Dynamica) to obtain plasma. Plasma protein and serum albumin were determined in 0.1 ml samples by the Biuret test (Smith et al., 1985) and the reaction with bromocresol green (cas 76-60-8), respectively. The commercial kits (Agat-Med) were used. The absorbance of the working solutions was measured at 540/625 nm against standards (spectrophotometer StatFax 303 Plus, Starmoff).

After the blood sampling, fish were euthanized with overdose of anesthetic. Samples of liver (distal lobe) and muscle (cut posterior to the dorsal fin) were collected and weighed with an accuracy of ±0.001 g (electronic balance GX-200, AND). The level of stored carbohydrates was analyzed in terms of liver glycogen content. The 2-mg fragment of liver was fixed in 96% ethanol and then digested in the laboratory in 500 µl 30% NaOH at 100°C for 30 min. Then, 100 µl of 60% H_2_SO_4_ was added to the sample, and the volume was brought up to 200 µl with Milli-Q water. The sample was mixed with 1.5 mg of 0.2% anthrone reagent (Sigma, cas 90-44-8) and incubated at 100°C for 20 min (Templeton, 1961). The optical density was read at 630 nm against glycogen standards.

Muscles were preserved in liquid nitrogen prior to lipid extraction. Lipids from 3.0-g samples of shredded muscle were extracted in methanol-chloroform mixture and purified with 1% KCl solution following Folch et al. (1957). The enzymatic hydrolysis reaction (Spinreact kit) provided the value of triglycerides (TAG, the main spare energy source); the indicator substance kinonimin was measured at 505 nm (Trinder, 1969).

To standardize the data on fatty acid (FA) processing status in the fish body, only six females of maximum size per morph were used to analyze the ratio of short chain (с < 20) saturated, long-chain (с ≥ 20) saturated, monoenic, unsaturated ω3 and ω6 unbound (free) FAs in samples of extracted lipids. The analysis was performed with the method of gas chromatography after FA methylation (Carreau and Dubacq, 1978). A Crystall-5000 (Chromateck) chromatograph with flame ionization detector was used (18 channels, phase width 0.25 µm, injector temperature 250⁰С, inlet pressure 1.25 MPa). A preliminary purification of methyl ethers of FAs was conducted with the method of thin-layer chromatography in benzol-hexane mixture. The FA groups were identified based on the comparative retention time and the values of the equivalent chain length (Ackman, 1969; Stransky et al., 1997).

### Data processing

The obtained data were statistically checked for the distribution matching using the Kolmogorov-Smirnov test (*p*>0.05 in all cases) and outliers were filtered based on the Chauvenet’s criterion. We found no significant physiological differences between males and females (t-test *p*>0.05 for all parameters, excluding FA data), nor was there a correlation between biochemical parameters and fish fork length in the combined sample (Spearman rank criterion < 0.35 *p*>0.05).

The effects of diet, habitat depth and morph identity on the parameter distributions were tested in a General Linear Model (GLM). ANOVA and a post-hoc Tukey HSD test were used to identify between-group differences; significance was considered at the *p*<0.05 level. To elucidate the overall difference in energy storage, we used Canonical Variate Analysis (CVA, based on Wilks’ Lambda statistics) of all measured parameters in six females of maximum size of all morphs. All calculations were performed in Statsoft v.10 (Hill and Lewicki, 2006).

## RESULTS

### Storage compounds

The fattening adult charrs possessed: 96–124 (mean 108.1) mg ml^-1^ proteins in plasma including 52–84 (66.6) mg ml^-1^ serum albumins; 56–142 (99.1) µg ml^-1^ glycogen in liver; and 17–43 (27.0) µg mg^-1^ TAGs in muscle. Using GLM (Table), we found a significant effect of morph identity on albumin and glycogen levels, and no effect of this predictor on plasma protein and TAG levels. Herewith, the levels of albumin, glycogen and TAGs were primarily statistically dependent on the diet of the charrs (insectivorous, amphipod-feeding, piscivorous, or deepwater benthos specialist). Muscle TAGs were also highly dependent on the depth of habitation (Table). The interaction of the feeding type and the depth of habitation effects significantly determined the content of albumins, glycogen and TAGs, and did not affect the content of plasma proteins.

**Table.**
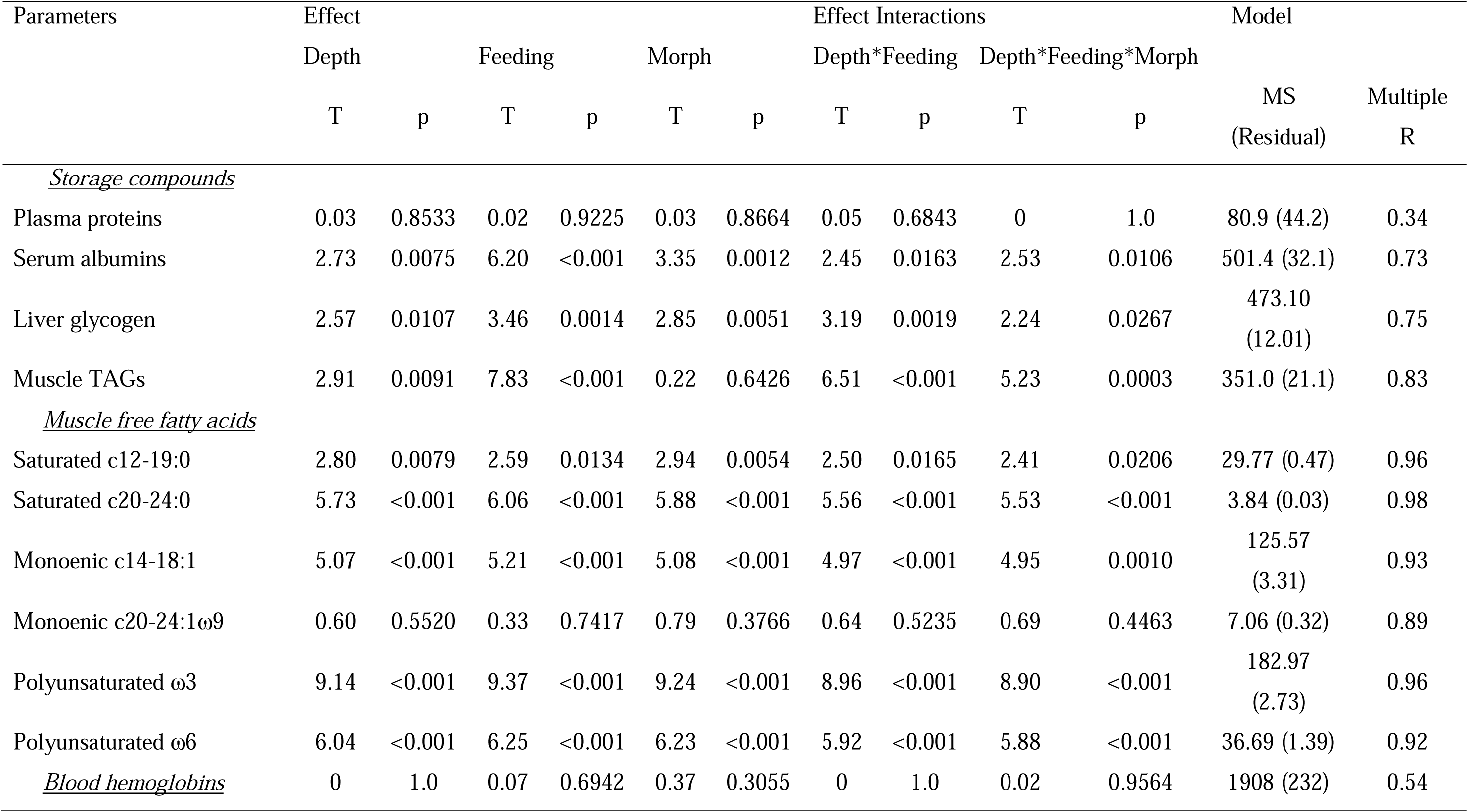
Statistical outputs from a General Linear Model for different biochemical parameters of eight morphs of Lake Kronotskoe charr.

Piscivorous morphs were characterized by high levels of serum albumins and muscle TAGs (especially L morph), and comparatively low level of carbohydrates in the liver. Conversely, insectivorous charrs differed in high liver glycogen and low serum albumins and muscle TAGs. Amphipod feeders had low levels of serum albumins and muscle TAGs, and the lowest level of liver glycogen. Deepwater consumer of silt benthos accumulated all types of spare substances (Fig. 2a–d). Using the Tukey HSD test, we found significant differences between the trophic groups for albumins (ANOVA F_7;148_ = 15.6 *p*=0.001), glycogen (F_7;127_ = 39.4 *p*<0.001) and TAGs (F_7;129_ = 16.7 *p*<0.001), and no significant difference for plasma proteins (F_7;134_ = 0.26 *p*=0.448). The latter parameter demonstrated the same trends of intergroup variation as serum albumins, but with a smaller range (concentration of albumins and total proteins in blood correlated, R = 0.78 *p*=0.001).

**Fig. 2.**
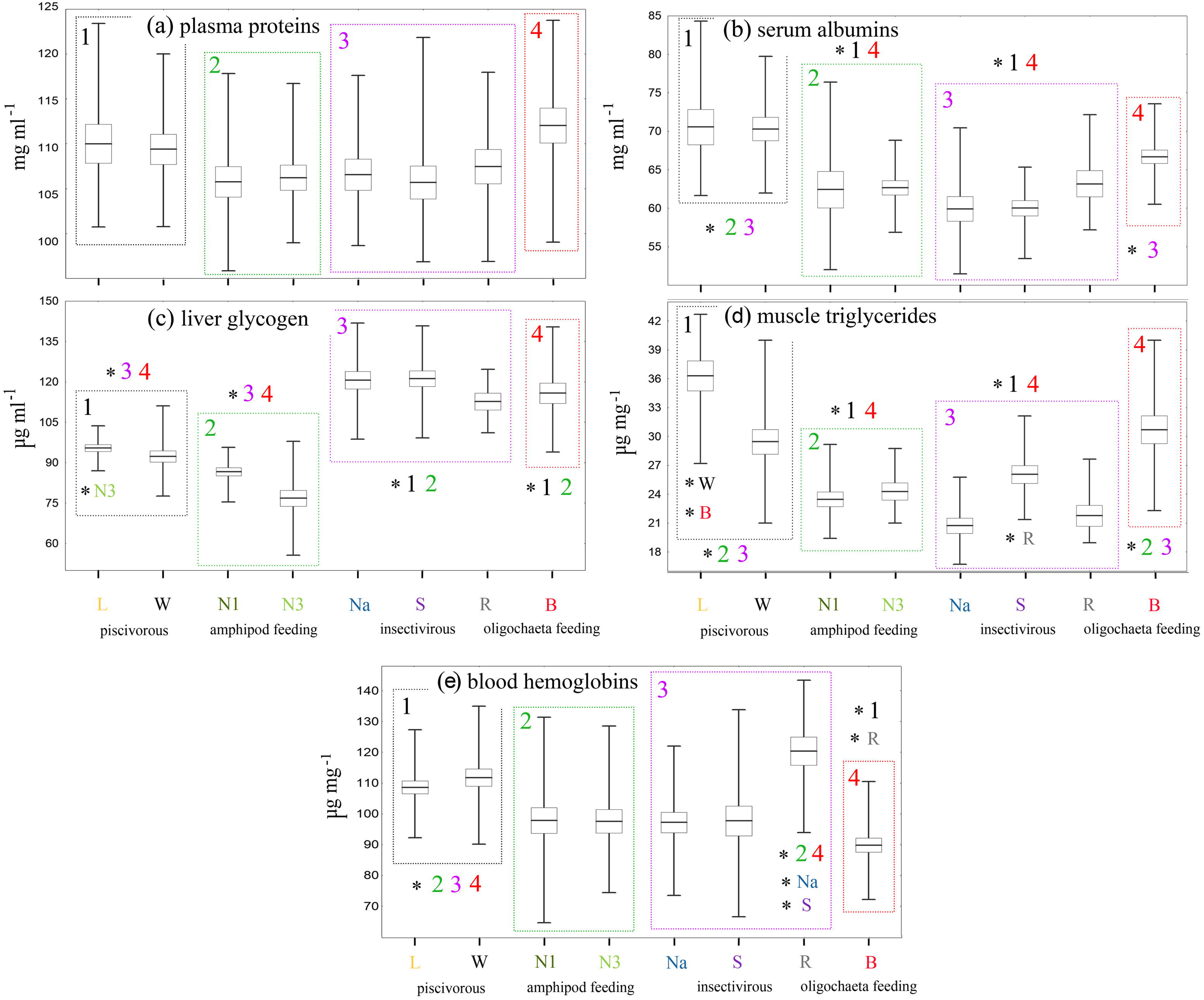
The content of plasma proteins (**a**), serum albumins (**b**), liver glycogen (**c**) and muscle TAGs (**d**), as well as blood hemoglobins (**e**) in eight morphs of Lake Kronotskoe charr. Mean (−) ± SE (□) and min−max values (|) are shown. Morphs are labeled; trophic groups are indicated by numbers: 1 − piscivorous, 2 − amphipod-feeding, 3 − insectivorous and 4 − oligochaeta (and mollusk)-feeding. Asterisks indicate trophic groups (morphs) for which significant differences (Tukey HSD *p*≤0.05) were found

### Fatty acids

The morph-specific ratio of the six FA groups is shown in Figure 3. Using GLM (Table), we found a significant influence of morph identity on the proportions of short-chain (с < 20) saturated, long-chain (с ≥ 20) saturated, short-chain monoenic, polyunsaturated ω3 and ω6 FAs. Herewith, the proportions of long-chain saturated, short-chain monoenic, ω3 and ω6 FAs were more dependent on the charr diet. The effect of depth was statistically more pronounced than that of diet for the proportion of short-chain saturated FAs (Table).

**Fig. 3.**
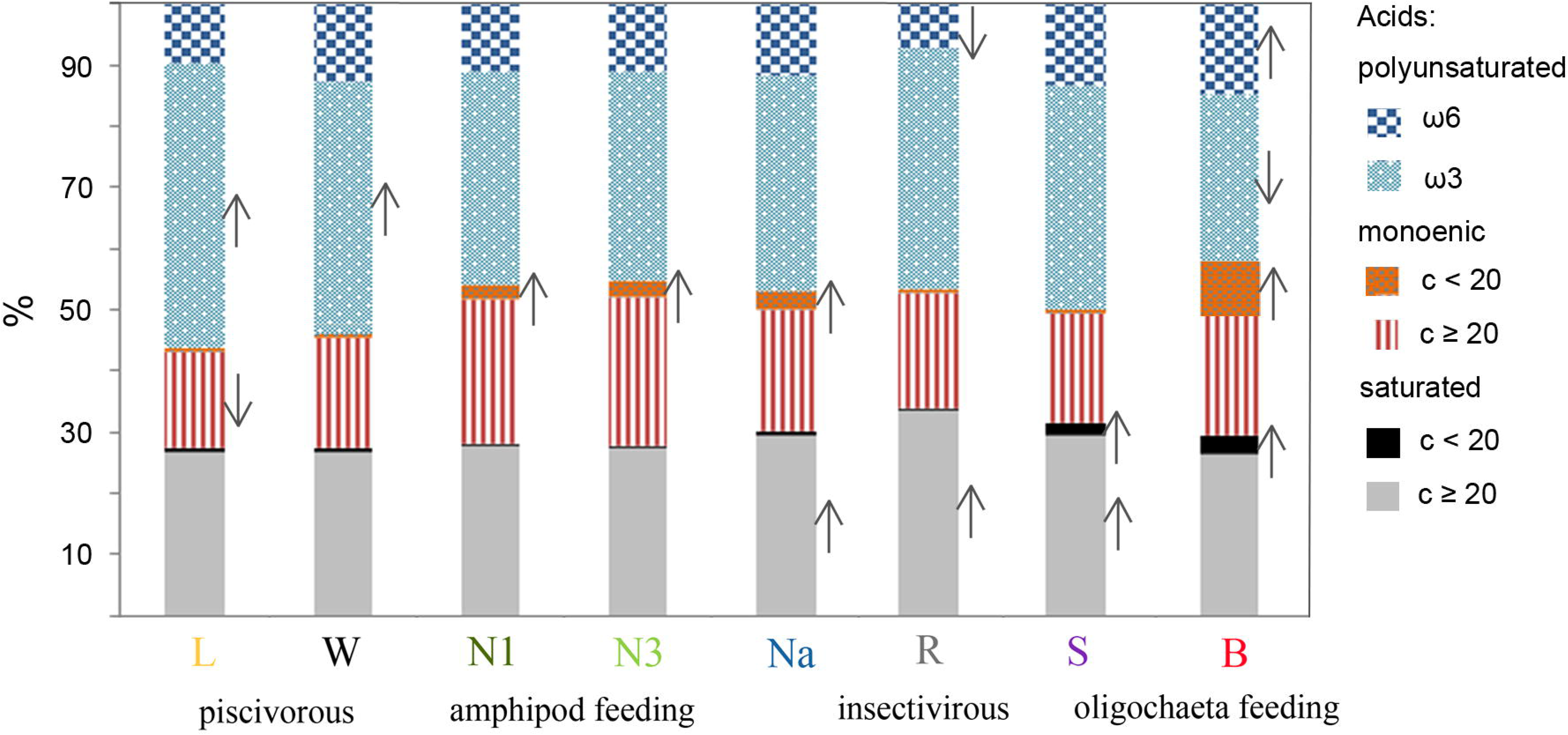
The ratio of free fatty acids with different chain types in the muscle of eight morphs of Lake Kronotskoe charr. Morphs are labeled; arrows indicate morphs that significantly differed (Tukey HSD *p*≤0.05) from others in the elevated or decreased ratio of the fatty acid group

Piscivorous morphs, especially L morph, were characterized by high levels of ω3 FAs. Insectivorous morphs differed in the selective accumulation of long-chain saturated FAs. The deepwater morph demonstrated significantly increased levels of long-chain monoenic ω9, ω6 and short-chain saturated FAs. The level of the latter FA group was also elevated in the semi-deepwater S morph. Three N morphs intensively accumulated short-chain monoenic FAs regardless of diet. R morphs showed a particularly low proportion of ω6 FAs (Fig. 3).

### Blood hemoglobins

The level of hemoglobins varied from 65 to 143 (mean 96.9) mg ml^-1^, and the intra-morph spread of the parameter reached 1.5−2 times. GLM failed to detect significant effects of diet, habitat depth and morph identity on the level of hemoglobins, MS (MS residual) = 1908 (232) and *p*=0.9564 (Table). However, using the Tukey HSD test (ANOVA F_7;146_ = 7.0 *p*=0.001), we were able to detect significantly elevated hemoglobin in the riverine R morph (*p*≤0.001) and both piscivorous morphs (*p*≤0.039) compared to the others. At the same time, the deepwater B morph had particularly low hemoglobin level (*p*<0.001; Fig. 2e).

### Physiological phenotypes

CVA of all measured biochemical parameters (except blood glucose) in six females of each morph allowed identification of morph-specific physiological phenotypes, F_77;181_ = 50.5 (*p*<0.001). The morphs occupied compact isolated areas in the CV1−CV2 space, and only areas of N1 and N3 morphs were brought closer together (Fig. 4a; eigen values are 149.9 for CV1 and 78.4 for CV2, while ≤ 35.0 for CV3−CV6). The individual classification probability of the model is 0.0128, canonical R = 0.958. The CV1 and CV2 axes discriminated morphs according to the feeding type and the depth of habitation, respectively.

**Fig. 4.**
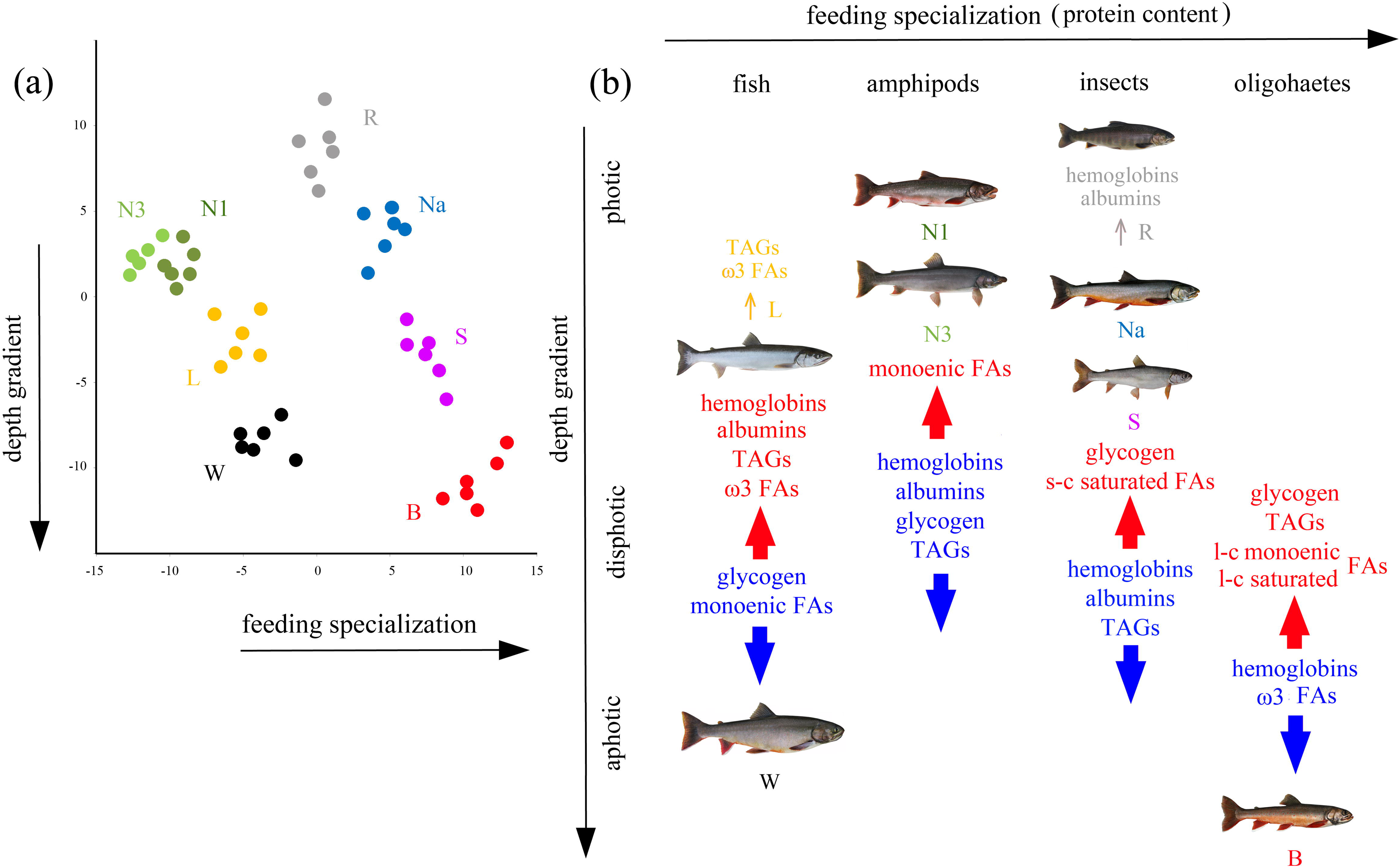
Results of physiological phenotype identification in eight morphs of Lake Kronotskoe charr. Canonical variate scaling of morphs (labeled) based on 11 biochemical parameters (**a**), and specifics of energy storage and metabolic rate depending on food composition and habitat depth (**b**). Red (blue) arrows indicate an increase (decrease) in component content compared to other morphs; additional small arrows indicate an increase in indicators with decreasing habitat depth

## DISCUSSUION

We found significant differences in the amounts and ratios of storage biopolymers among ecomorphs that diverged from the common ancestor Dolly Varden within a few thousand years in a single ecosystem. Piscivorous morphs maintain high levels of hemoglobin and store lipids, including polyunsaturated FAs, the ability to process proteins for energy, and elevated levels of metabolic rate indicators. The insectivorous morphs differ from each other in proxy indicators of metabolic rate, but have a high content of carbohydrates, as well as low albumin and TAG levels. The amphipod consumers are characterized by low carbohydrate, albumin and TAG levels. The metabolic characteristics of the deepwater benthivorous feeder do not resemble those of the other groups. This ecomorph does not maintain high level of hemoglobin, but accumulates spare substances. Each ecomorph can be identified based on multivariate analysis of measured physiological parameters, i.e. it has a specific metabolic phenotype that tightly correlates with lifestyle (Fig. 4). Our findings support the hypothesis that the metabolic phenotype is under selection during the adaptive radiation of Lake Kronotskoe charrs.

Physiological adaptations ensure vital homeostasis and fitness under current external conditions (Koedijk et al., 2010; Seebacher et al., 2015; Cloyed et al., 2019), and are shaped by both the influence of habitat surrounding/feeding strategy and inherited characteristics (Cui and Liu, 1990; Soengas et al., 2007; Goetz et al., 2013; Chavarie et al., 2016). Our data clearly demonstrate that the diversity of metabolic characteristics of the Lake Kronotskoe charrs, in particular differences in the amount of reserved substances correlate well with the trophic specialization of ecomorphs. Food objects differ in the proportion of lipids, peptides and carbohydrates: salmonid flesh contents of 7–8% lipids, 20% protein and 1% carbohydrate (Pozdnyakova et al., 2018); gammarids – 25–30%, 40% and 6% (Mathias et al., 1982); chironomid larvae – 15%, 55% and 20%; and oligochaetes – 10–15%, 60% and 7–10% (De La Noue and Choubert, 1985), respectively. Such discrepancies in food composition are consistent with the maximum carbohydrate values in the liver of insectivorous morphs, the low carbohydrate concentration in piscivorous and amphipod consumers, and the high plasma protein and albumin content in oligochaete specialist. Herewith, food composition do not explain the low TAG levels in amphipod feeders and the highest TAG levels (and relatively high plasma protein) in piscivorous morphs, which are likely determined by internal factors.

Ecomorphs also differ in the composition of FAs, the dominant source of metabolic energy in cold-water fish (Tocher et al., 2003). As expected, piscivorous charrs display a maximum content of ω3 FAs, which are re-deposited in fish feeders (Sushchik et al., 2006; Chavarie et al., 2016) and are essential compounds for maintaining rapid catabolism (Sargent et al., 2002; Schmitz and Ecker, 2008; Tallima and El Ridi, 2018). The amphipod specialists (N1 and N3) accumulate more monoenic FAs, whereas the insectivorous morphs have an elevated ratio of saturated FAs. Similar differences in FA content have been reported for amphipod- and insect-feeding morphs in the *S. malma* population from Lake Dal’nee (Busarova et al., 2017a). The demersal oligochaete consumer displays a decreased level of ω3, but an elevated level of saturated, polyunsaturated ω6 and monoenic ω9 FAs. Since monoenic ω9 FAs are transmitted by pelagic crustaceans (Iverson, 2009), their high content in deepwater demersal fish is quite surprising. We assumed that such a deviation in the composition of FAs has an intrinsic background, since the FAs ratio in salmonids is determined not only by diet (Happel et al., 2016; Gladyshev et al., 2017), but also by genetics (Horn et al., 2018; Gladyshev et al., 2022a; 2022b).

The lipid content (TAGs and free saturated FAs) in muscle is associated with the depth of habitation. Such a positive correlation could be an adaptive response to temperature decrease, as enzymes of lipid oxidation lose activity more slowly with a decrease in temperature than enzymes of carbohydrate oxidation (Crockett and Sidell, 1990). The pelagic morphs probably accumulate an additional amount of lipids to increase valuable buoyancy (Phleger, 1998; Goetz et al., 2013). During the juvenile stage, S morph is distributed in the water column, and is characterized by the highest lipid content, whereas the piscivorous L morph inhabiting the lake tributaries is at the stage of faster growth and is distinguished by the lowest lipid content (Esin et al., 2018). After the lake migration and switch to piscivory, L morph begins to actively accumulate lipids.

Ecomorphs display significant differences in hemoglobin level, an adaptive trait that varies depending on lifestyle, locomotor activity and habitat conditions (Hall and Gray, 1929; Zaprudnova et al., 2015; Esin et al., 2021). High hemoglobin in the blood of the riverine and piscivorous charrs is probably an adaptation that ensures an increased swimming activity. They also have comparatively elevated serum albumin and total plasma protein levels. The low hemoglobin and plasma proteins level inherent in ecomorphs with a moderate lifestyle, with the exception of B morph, which has a high level of plasma proteins and albumin.

Differences in indicators of metabolic activity: the level of hemoglobin and plasma proteins, along with previously revealed differences in the content of thyroid hormones (Markevich et al., 2021), the crucial regulators of fish metabolic activity (Blanton and Specker, 2007; Deal and Volkoff, 2020; Zwahlen et al., 2024), allow suggesting that the metabolic rate decreases in the series ‘L - W - R / B - N3 - N1 - Na - S’. The high metabolic rate probably allows piscivorous morphs (L-W) to accumulate lipids as a reserve, even though the fish they consume are less fatty than amphipods and insects. Such processing should be physiologically beneficial because lipids differ in a high energy content (∼39 J mg^-1^) compared to peptides (∼24 J mg^-1^) and carbohydrates (∼17 J mg^-1^) (Cavaletto and Gardner, 1999). The amphipod feeders require an intensification of energy-intensive digestive processes for a more efficient assimilation of chitin-reach amphipod bodies, while consumers of more easily digestible insects (Na-S-R) shift metabolism towards carbohydrate accumulation at the expense of expenditure on somatic growth. The deepwater oligochaete consumer (B) displays the most specific values of biochemical parameters. It has the lowest hemoglobin level, but high plasma proteins and albumin level, accumulates both glycogen and TAGs, and uses relatively poorly oxidized monoenic and saturated fatty acids for energy.

Thus, each ecomorph possesses a specific metabolic phenotype corresponding to its tropho-ecological specialization and lifestyle. Given that many ecomorph-specific traits, such as the low content of ω6 FAs in the riverine ecomorph; the accumulation of short-chain monoenic FAs by monophyletic, but trophically distinct N ecomorphs; or the high level of ω9 FAs in the deepwater ecomorph, cannot be explained by differences in diet and habitat per se, we assume an intrinsic, probably genetic background of the physiological diversity of charrs. Similar genetically based metabolic differences have been reported for the various wild and farmed salmonids, including Arctic charr, a phylogenetically closed species to *S. malma* that demonstrates numerous examples of lacustrine adaptive radiation (Horn et al., 2018; Gladyshev et al., 2022a; 2022b).

The metabolic phenotype is a very complex and polygenic trait, the Lake Kronotskoe charr assemblage is very young, and the probability of emergence and accumulation of multiple genetic discrepancies among ecomorphs seems elusive. Under these circumstances, we suggest specific selection on several downstream regulators of energy metabolism, among which endocrine signals: thyroid hormones and leptin are the most plausible candidates. Our assumption is based on knowledge of the physiological and ecological functions of these closely related hormones. They play a crucial role in the regulation of the citric acid cycle and lipid metabolism, as well as in the transformation of energy metabolism in response to environmental changes and during the ontogenetic stage transition, accompanied by a shift in trophic and habitat specialization (Zimmermann-Belsing et al., 2003; Deck et al., 2017; Weidner et al., 2020; Blanco and Soengas, 2021; Martelly and Brooks, 2023; Zwahlen et al., 2024). Our hypothesis is also supported by data on the involvement of the thyroid axis in the lacustrine and riverine adaptive radiation of *Salvelinus* (Esin et al., 2021a,b), differences in thyroid hormone content among ecomorphs (Markevich et al., 2021) and selection on thyroid and leptin genes during the adaptive radiation of the Lake Kronotskoe charrs (Woronowicz et al., 2023). Taking together, these data indicate that physiological phenotype, particularly the endocrine control of energy metabolism, is an important target of selection during adaptive radiation. Furthermore, given that changes in most, if not all, environmental factors primarily affect the energy balance of the organism, and that changes in physiology are required to maintain a state of energy equilibrium, we suggest that the metabolic phenotype is the primary biological characteristic that should increase variance and shift the mean of fitness in a way that increases niche availability when the ancestral lineage faces an ecological opportunity. Such physiological shifts affect not only energy metabolism, but also development, behaviour and life strategy, thus orchestrating complex adaptive transformations of diversifying lineages.

## Acknowledgments

The authors would like to express their deep gratitude to Dr Dmitriy V. Zlenko and Dr Nikolay O. Melnik, who participated in the work at Lake Kronotskoe. We are also grateful to Dr. Daria M. Panicheva for organizing the logistics of the expeditions. The study was conducted in frame of the IEE RAS research program no. 0089-2021-0003 and the VNIRO state assignment no. 076-0007-22-00. The research is supported by the academic leadership program “Prioritet-2030” of the Ministry of Science and Higher Education of Russia (national project “Science and universities”).

## Ethics statement

All procedures with fish were carried out according to the guidelines and following the laws and ethics of the Russian Federation, and approved by the ethics committee of the Severtsov Institute of Ecology and Evolution, Russian Academy of Sciences (Approval ID: N 88 issued on 22.03.2024). Fish was sampled in accordance with Permission № 02/2022 issued on 23/06/2022 by the Ministry of Natural Resources and Ecology of the Kamchatka Krai.

## Author Contributions

Evgeny Esin and Grigorii Markevich conceived the ideas and designed methodology; Grigorii Markevich, Elena Shulgina and Fedor Shkil collected the data; Yulia Baskakova and Roman Artemov processed the samples; Evgeny Esin and Elena Shulgina analyzed the data; Evgeny Esin and Fedor Shkil led the writing of the manuscript. All authors contributed critically to the drafts and gave final approval for publication.

